# Functional Near-Infrared Spectroscopy (fNIRS) informed neurofeedback: regional-specific modulation of lateral orbitofrontal activation and cognitive flexibility

**DOI:** 10.1101/511824

**Authors:** Keshuang Li, Yihan Jiang, Yilong Gong, Weihua Zhao, Zhiying Zhao, Xiaolong Liu, Keith M. Kendrick, Chaozhe Zhu, Benjamin Becker

## Abstract

Cognitive flexibility and reward processing critically rely on the orbitofrontal cortex. Dysregulations in these domains and orbitofrontal activation have been reported in major psychiatric disorders. Haemodynamic brain imaging informed neurofeedback allows regional-specific control over brain activation and thus may represent an innovative intervention to regulate orbitofrontal dysfunctions. Against this background the present proof-of-concept study evaluated the feasibility and behavioral relevance of functional Near-Infrared Spectroscopy (fNIRS) assisted neurofeedback training of the lateral orbitofrontal cortex (lOFC). In a randomized sham-controlled between-subject design 60 healthy participants underwent four subsequent runs of training to enhance lOFC activation. Training-induced changes in the lOFC, attentional set shifting performance and reward experience served as primary outcomes. Feedback from the target channel significantly increased regional-specific lOFC activation over the four training runs in comparison with sham feedback. The experimental group demonstrated a trend for faster responses during set shifting relative to the sham group. Within the experimental group stronger training-induced lOFC increases were associated with higher reward experience. The present results demonstrate that fNIRS-informed neurofeedback allows regional-specific regulation of lOFC activation and may have the potential to modulate associated behavioral domains. As such fNIRS-informed neurofeedback may represent a promising strategy to regulate OFC dysfunctions in psychiatric disorders.

## Introduction

Neurofeedback techniques have gained increasing attention as non-invasive means to regulate brain function for scientific and therapeutic purpose [1,2]. The corresponding methods employ a biofeedback approach that uses real-time information of brain activity to enable self-regulation of a particular neural signal [1,3]. Compared with the traditionally used Electroencephalography (EEG) neurofeedback, the use of haemodynamic imaging signals as near real-time neural feedback is relatively new. The majority of haemodynamic neurofeedback studies employed functional magnetic resonance imaging techniques [1,3] and a growing number of studies have demonstrated that functional magnetic resonance imaging (fMRI) neurofeedback-guided regulation of regional brain activity can produce changes in cognitive and emotional processes specifically associated with the target brain region (e.g. [4]; overview see [1,2]). Based on accumulating evidence for the behavioural relevance of the neurofeedback-induced modulation of brain function, initial studies evaluated the potential of fMRI-neurofeedback as promising innovative intervention for psychiatric disorders [5].

Although the therapeutic potential of fMRI-guided neurofeedback has been documented in initial randomized controlled trials [6,7], translation into clinical applications is hampered by the high costs of MRI assessments, the rather stressful MRI environment as well as limitations inherent to the MRI method, particularly a high sensitivity for motion and physiological noise [8], and field inhomogeneities caused by different magnetic susceptibility to air and tissue resulting in signal loss in orbitofrontal regions [9]. A previous fMRI neurofeedback study demonstrated that optimized MRI imaging parameters can improve signal quality in the orbitofrontal cortex (OFC) [10], however, imaging orbitofrontal regions-particularly under real-time signal processing conditions - remains challenging.

Functional near-infrared spectroscopy (fNIRS) is a non-invasive optical neuroimaging technique which – similar to blood-oxygen-level dependent (BOLD) MR imaging – can be employed to detect changes in haemoglobin concentration associated with neural activity [11]. Briefly, neural metabolism is supported through a localized vascular response that causes an influx of oxygen-rich blood to the active region, reflected by a regional increase in oxy-haemoglobin (oxy-Hb) and a decrease in deoxy-haemoglobin (deoxy-Hb) [12]. fNIRS measures the concentration of oxygenated haemoglobin (oxy-Hb) and deoxygenated haemoglobin (deoxy-Hb) in cerebral vessels according to their absorption spectra for light in the near-infrared range [13]. Despite limitations, particularly a restricted penetration depth and resolution, fNIRS has been increasingly employed in cognitive and clinical neuroscience. Due to recent technological and methodological progress in fNIRS imaging and advantages over fMRI, including lower costs, easy application and robustness against motion and susceptibility artefacts, fNIRS has become an attractive hemodynamic imaging alternative.

The OFC, a ventral subdivision of the prefrontal cortex, is cytoarchitectonically and functionally heterogenous region with dense connections to cortical and subcortical areas including sensory regions as well as limbic and striatal regions [14]. Supported by the subregion-specific segregated circuits the OFC contributes to several highly integrative functional domains including emotion, decision making, value coding and behavioural flexibility [15,16]. Across species lesions of the OFC produce marked impairments in reversal learning and attentional set shifting, suggesting a critical role of the OFC in cognitive flexibility [15]. Studies employing reversal learning paradigms reported that rodents with OFC damage could learn the initial discrimination by responding to one cue to receive reward and to withhold or inhibit a response to avoid punishment or non-reward. However, after cue-outcome associations are reversed, OFC-lesioned rodents required considerably longer to adapt their behaviour [17-22]. More recently the role of the OFC in reversal learning has additionally been confirmed by human fMRI studies. These studies used rewarding and punishing stimuli, and reported that the adaptation to changing reinforcement contingencies was mediated by the OFC. Cognitive flexibility additionally encompasses attentional set-shifting supporting flexible behavioural adaptation in the context of relevant and irrelevant information. Set shifting paradigms have been combined with fMRI to dissect regional-specific contributions of the prefrontal cortex to cognitive flexibility subdomains and it has been reported that the attentional control sub-facet engages the ventrolateral cortex whereas reversals specifically engage the lateral orbitofrontal cortex [23]. In addition to cognitive flexibility, the role of the OFC in reward processing, specifically modulating outcome expectancies related to reward, has been well documented. For instance, human neuroimaging studies implicated the OFC in the anticipation and evaluation of expected outcomes [24] as well as value-guided decision making [25].

In line with the important role of the OFC in cognitive flexibility and reward processing, neural alterations in this region have been consistently reported in psychiatric disorders characterized by dysregulations in these domains, most prominently addictive disorders, obsessive compulsive disorder [15], attention-deficit/hyperactivity disorder [26] and major depression [27]. Given that the efficacy of the established pharmacological and behavioural treatment options for these disorders is limited [28], it has been advocated to employ hemodynamic imaging informed neurofeedback to modulate the identified neural alterations in a regional-specific manner to alleviate psychiatric symptoms and normalize functional deficits [28-31].

To facilitate translation of haemodynamic neurofeedback into clinical applications, the present randomized, sham-controlled study aimed at evaluating the feasibility to employ fNIRS-informed neurofeedback as strategy to modulate brain activity. Initial studies have demonstrated the feasibility to use fNIRS-informed neurofeedback to allow participants to acquire regulatory control over brain activity in motor related regions [32,33]. Subsequent studies demonstrated the feasibility to gain regulatory control over prefrontal brain activity via fNIRS-informed neurofeedback and its potential to enhance executive functions [34]. Given the important contribution of OFC dysregulations to the functional impairment in psychiatric patients as well as the methodological and practical challenges to use fMRI-based neurofeedback approaches in this context, the present fNIRS neurofeedback study targeted the lateral OFC. Importantly, previous studies have shown that fNIRS can sensitively detect deoxyhaemoglobin changes in the OFC [35], and reliably assess neural activity in the OFC during cognitive flexibility and reward evaluation [36]. Moreover, an increasing number of studies employed fNIRS imaging to associate aberrant cognitive flexibility and reward processing with altered OFC activation in psychiatric patients [37]. To further determine the functional relevance of the neurofeedback induced OFC activation changes, behavioural indices of cognitive flexibility and reward processing – both of which have been associated with the lateral OFC [15] - served as primary behavioural outcomes to evaluate the training success. To control for unspecific effects of training, the fNIRS-informed OFC neurofeedback was embedded in a randomized sham-controlled between-subject experiment with a total of n = 60 health participants. Based on previous studies [4,6], we expected that participants in the experimental but not the sham feedback group would learn to successfully up-regulate regional-specific activity in the lateral OFC and that this would be accompanied by increased cognitive flexibility and reward experience on the behavioural level.

## Methods and Materials

### Participants

N = 60 healthy young males were enrolled in the present study. The main aim of the study was to determine the feasibility and functional relevance of real-time fNIRS-informed neurofeedback training targeting the lateral OFC. To reduce variance related to sex-differences or effects of menstrual cycle on OFC activity and the primary behavioural outcomes [38], the present proof-of-concept experiment focused on male participants. To control for unspecific effects of the training procedures on the primary outcomes the neurofeedback training was embedded in a randomized, sham-controlled between-subject experimental design. Participants were randomly assigned (30 participants in each group) to receive either real-time feedback from the OFC target region (experimental group) or a sham feedback (control group) during the training. Participants were randomized without stratifying for further variables. All participants provided written informed consent. The study had full ethical approval by the local ethics committee of the University of Electronic Science and Technology of China and the procedures were in accordance with the latest version of the declaration of Helsinki.

### Neurofeedback training protocols and procedures

The training session included four subsequent runs of alternating rests and regulation blocks (four blocks per run, block duration 25s). The experimental group received real-time feedback from fNIRS channel No.7 located over the right lateral OFC (location of the optodes and feedback channel are displayed in Figure 1), whereas the control group received feedback from a participant who had previously underwent the experimental training (similar approach in [32]). To accustom the participants to the equipment and to reduce variance related to trial-and-error attempts during the initial training blocks the training was preceded by a pre-training session during which all subjects received OFC feedback and were required to explore a suitable regulation strategies during 6-10 feedback blocks. When participants reported that they had discovered an effective strategy a recovery break of 15-30min was included before the training session started. Participants were asked to employ the strategies they discovered during the pre-training session to increase lateral OFC activity during the training. To motivate participants, the feedback was displayed as a stone-lifting game (protocols and feedback displayed in Figure2). Briefly, the higher the stone floats above the ground the higher is the neural activity as measured by the brain signal in the chosen feedback channel No.7.

**Figure 1.**
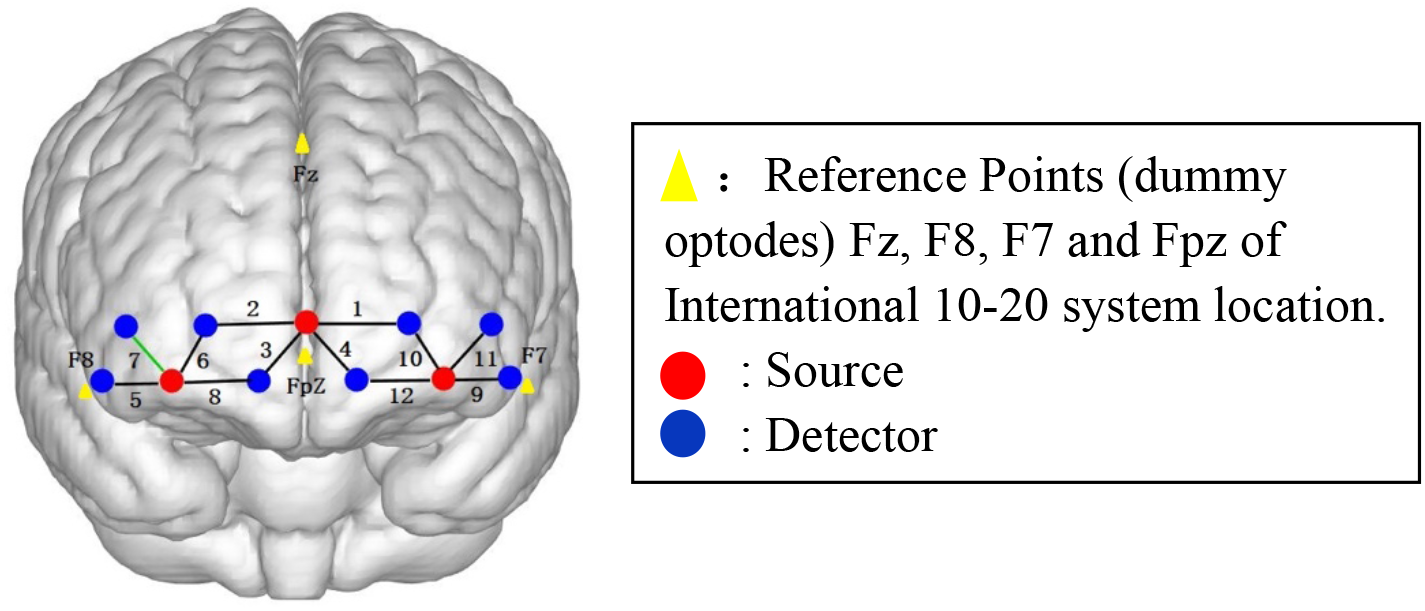
Optodes configuration (3 sources: red dots, 8 detectors: blue dots). Each source and each detector were 30 mm apart. A black line connecting a detector and a source represents a measurement channel and its number. 6 channels were used per hemisphere: 2 channels covering the medial OFC (right hemisphere: channels 2 and 3; left hemisphere: channels 1 and 4) and, 4 channels covering the lateral OFC (right hemisphere: channels 5-8; left hemisphere: channels 9–12). The probe was projected on the scalp with the anchor points (visualized as yellow triangles) corresponding to Fz, F7, and F8 in the International 10–20 system. The source in the middle of the probe array was located on the Fpz. Optode placement was anatomical symmetrical in both hemispheres. Channel No.7 (visualized as green line) served as target channel during the neurofeedback training.

**Figure 2.**
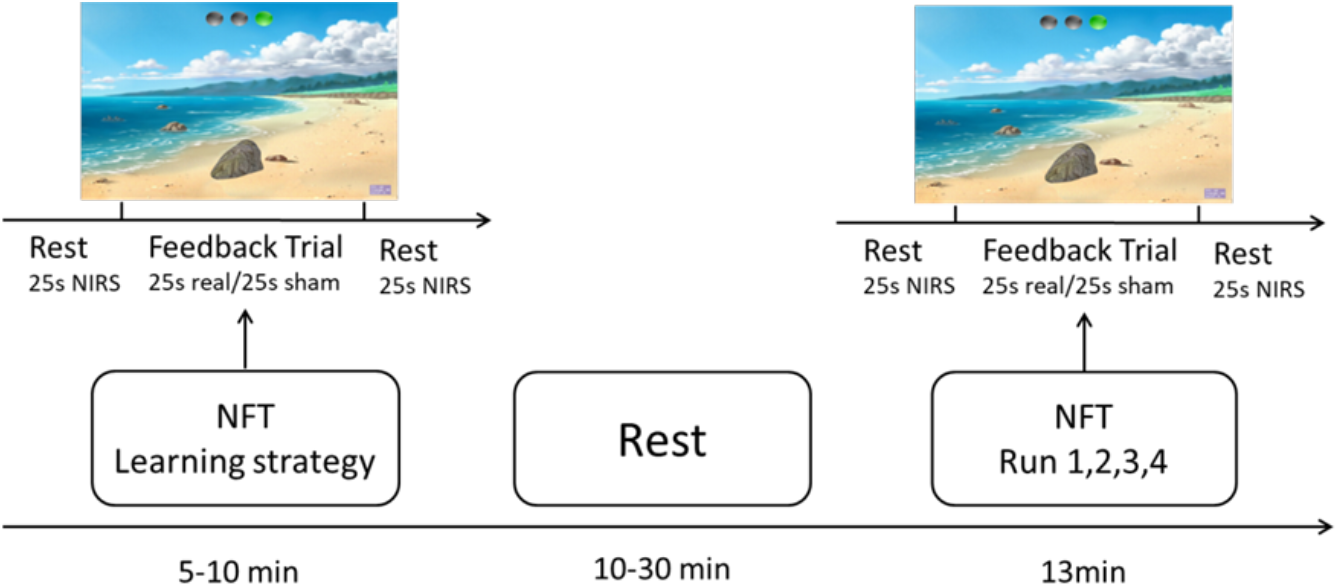
Experiment procedures for two training sessions.

The condition (rest / regulate) is visually indicated to the participant via 3 lights on the top. At the beginning, the red light on the left side is on, which instructs the participant to rest, and after that it switches into the green light on the right side which instructs the participant to lift the stone presented on the screen. The participants were asked to try to lift the stone as high as possible by regulating their brain activity. Participants were informed that the purpose of the training was to test whether they could learn to up-regulate their OFC activity. To increase their regulation ability the neural regulation success would be visually presented to them via the stone lifting game (all subjects were explicitly informed that the higher the stone floats the higher is the OFC regulation success). Given that an explicit strategy instruction is not necessary for successful neurofeedback-assisted acquisition of neural regulation [39,40], no explicit strategies for regulation were provided to the participants. Participants were instructed not to control the stone by physical means such as breathing or head/body motion but rather to discovery efficient mental control strategies. Once they discovered an efficient strategy to lift the stone during the pre-training they were asked to continue using the strategy during the subsequent training sessions.

To determine the functional relevance of the training on the behavioural level, participants were administered the Intra-Extra Dimensional Set Shift (IED) task and rated their rewarding experience on a visual analogue scale (VAS) after the training session. The IED paradigm has been widely employed to examine cognitive flexibility and has a high sensitivity for changes in fronto-striatal functioning [41]. In this task, participants are required to use the provided feedback to discover a rule that determines which stimulus is correct. After six correct responses, the stimuli and/or rule changes. Initially the task involves simple stimuli (two different pink shapes), during this stage shifts in the rule are intra-dimensional. During alter stages of the task compound stimuli are used (e.g. white lines overlay the pink shapes, corresponding shifts refer to extra-dimensional shifting. To assess effects of training on reward experience participants were additionally asked to rate their liking of the training on a 1-9 (“How much did you enjoy the task”) scale.

### NIRS Data acquisition, online pre-processing and neurofeedback

Haemodynamic response (HR) signals were assessed using one NIRSport fNIRS Systems (8 sources/8 detectors, NIRx Medizintechnik, GmbH, Berlin, Germany) coupled in tandem-mode and operating at two wavelengths (760 and 850 nm) at a sampling rate of 20.83 Hz. An optode-set of 3 sources and 8 detectors was used leading to 12 source detector pairs (channels; see Figure 1) [optode placement in line with 42].

Online pre-processing of the NIRS signal was performed by the built-in real-time output solution implemented in the NIRSport system. Next, the real-time output was computed and visually displayed by via a previously evaluated real-time fNIRS neurofeedback platform [43]. As a first step, the raw oxy-Hb NIRS signal was smoothed using a 2s moving average window. A baseline was calculated by taking the average of signals 2s before each regulation block and was subsequently subtracted from the smoothed signal. For each participant, the coefficient of difficulty is individually adopted based on the individual maximum activation intensity as determined by a preliminary test. The feedback signal was normalized to the same difficult by dividing this coefficient. Next, changes in the feedback signal during the regulation period were provided as visual feedback on a screen. Participants in the sham neurofeedback group received oxy-Hb signal changes from a participant in the real-time OFC neurofeedback group who had previously completed the training. Participants were blinded for the training condition they received and the experimenter operating the feedback platform was separated from the participant by a partition wall to minimize interaction during pre-training and training.

### Offline preprocessing and analyses

The fNIRS raw data were pre-processed and analysed using the NIRS toolbox in SPM (Statistical Parametric Mapping, https://www.fil.ion.ucl.ac.uk/spm/)[44] and in-house scripts in MATLAB (The MathWorks, Inc.). During pre-processing, a second-order detrending was applied to remove baseline drifts and low-pass filtering (Gaussian smoothing with Full width at half maximum (FWHM) 4 sec) was employed to remove high-frequency noise. The subsequent data analysis focused on the oxyhemoglobin (HbO) signal as it has been demonstrated to exhibit larger signal changes and higher sensitivity to task-related neural activation compared to the deoxyhemoglobin signal [45-47]. A generalised linear model (GLM) approach was employed to model the task-related hemodynamic response on the individual level. Beta estimates were obtained for each participant and channel. The primary outcome to determine training success on the neural level were HR activity changes over the training runs in the target channel (right lateral OFC; channel No.7). To further control for unspecific effects of training or effects of mental effort on OFC activity, individual-level beta-values from the target channel were subjected to group-level activation analyses comparing the experimental and the control group. Differences were considered significant using a channel-level threshold of p < 0.05 (FDR corrected).

### Primary outcomes and evaluation of training success

Training-induced changes in the OFC target channel served as primary outcome to evaluate the training success on the neural level. Channel specific activity beta-estimates were employed as dependent variable and effects of training were determined by means of mixed ANOVA models including the between-subject factor group (experimental vs sham control) and the within-subject factor training run (run1/run2/run3/run4). Significant effects were further explored by means of appropriate Bonferroni-corrected post-hoc tests. To evaluate the functional relevance of training on the behavioural level post-training IED performance indices and liking ratings were compared between the two training groups. Associations between neural and behavioural training success were explored by means of analysing correlations between training-induced lOFC changes (run4 > run1) and behavioural indices within the training groups. Statistical analyses were carried out using SPSS version 22 (IBM, Inc).

### Up-regulation strategies

After the training sessions, all participants were asked to report the strategies they employed during the training. The reported strategies were qualitatively assessed by six independent male raters. To control for different strategies between the training groups the frequencies of the reported contents was compared between the experimental and sham group using Pearson x ^2^ test [48].

### Control of potential confounders

To further control for confounding effects of pre-training between-group differences in mood and psychopathological symptom load, corresponding indices were assessed by means of the PANAS (the Positive and Negative Affect Schedule [49]), SAI (State Anxiety Inventory [50]), BDI II (Beck Depression Inventory-II [51]) and BIS (Barratt Impulsiveness Scale [52]) administered before the training sessions. To further control for confounding effects of between-group differences in the perceived training success all participants were required to rate their training success (scale ranging from −4 to 4).

## Results

### Data quality control

Two participants reported that they were not able to gain control over their brain activity in the pre-training test runs and were thus excluded from the subsequent neurofeedback training. Initial examination of data quality identified one participant from each group as an outlier with respect to data from the training channel during at least two runs (>two standard deviations from the group mean at a given assessment, additionally confirmed by the SPSS outlier detection function). Consequently, data from these participants were excluded from all analysis resulting in a total of n = 56 participants for the primary analysis (n = 27, experimental group; n = 29 control group).

### Mood states and other psychological conditions

The training groups did not differ with respect to age (M_experimental_ = 21.3 years, SD = 2.02; M_control_ = 21.7 years, SD = 2.08; t_df_ =-0.656, p = 0.515) or pre-training mood and psychopathological load (Table1, t_df_ < 1.463, ps > 0.149). Moreover, the training groups reported a comparable evaluation of their perceived training success (t_df_ =1.009, p = 0.317), arguing against confounding effects of between-group differences in the experienced success during training.

**Table 1.**
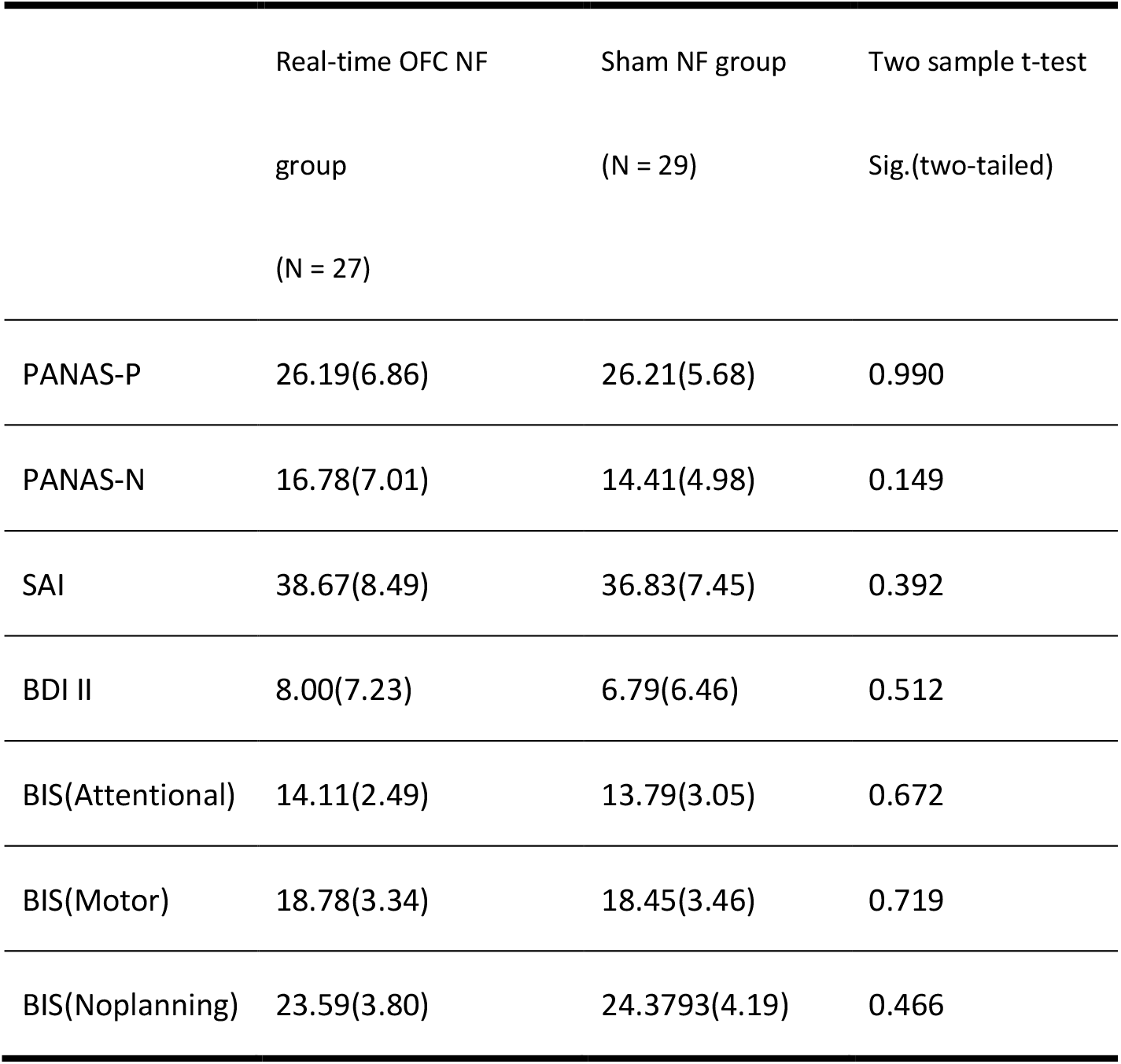
Pre-training mood and psychopathological symptom load in the two training groups, mean and SDs (in brackets) are reported. Abbreviations: PANAS-P, the Positive and Negative Affect Schedule – positive; PANAS-N, the positive and negative affect schedule – negative; SAI, State Anxiety Inventory; BDI II, Beck Depression Inventory-II); BIS, Barratt Impulsiveness Scale.

### Evaluation of training success – primary neural outcome

A mixed two-way ANOVA with the factors run (run1/run2/run3/run4) and group (real feedback vs. sham feedback) and the dependent variable lateral OFC activity as measured by the beta-values from the target channel revealed a main effect of run (F_(3,162)_ = 7.51, p = 0.001) and group (F_(1,54)_ = 15.36, p < 0.001) as well as a run*group interaction effect (F_(3,162)_ = 5.32, p = 0.005). Post-hoc comparisons demonstrated that activation in the target channel significantly increased over the course of the real feedback training (Run1 < Run3, p < 0.001; Run1 < Run4, p = 0.002; Run2 < Run3, p < 0.001; Run2 < Run4, p = 0.013, two-tailed). Concordant analysis of the sham training data did not yield significant changes in the target channel (all ps > 0.151, two-tailed). Directly comparing the training groups further revealed that the experimental group exhibited significantly higher activity during runs 3 and 4 (p < 0.001, two-tailed) in the lateral OFC target channel as compared to the control group, however, not during runs 1 and 2 (p > 0.08, two-tailed, see Figure 3) further confirming training success on the neural level.

**Figure 3.**
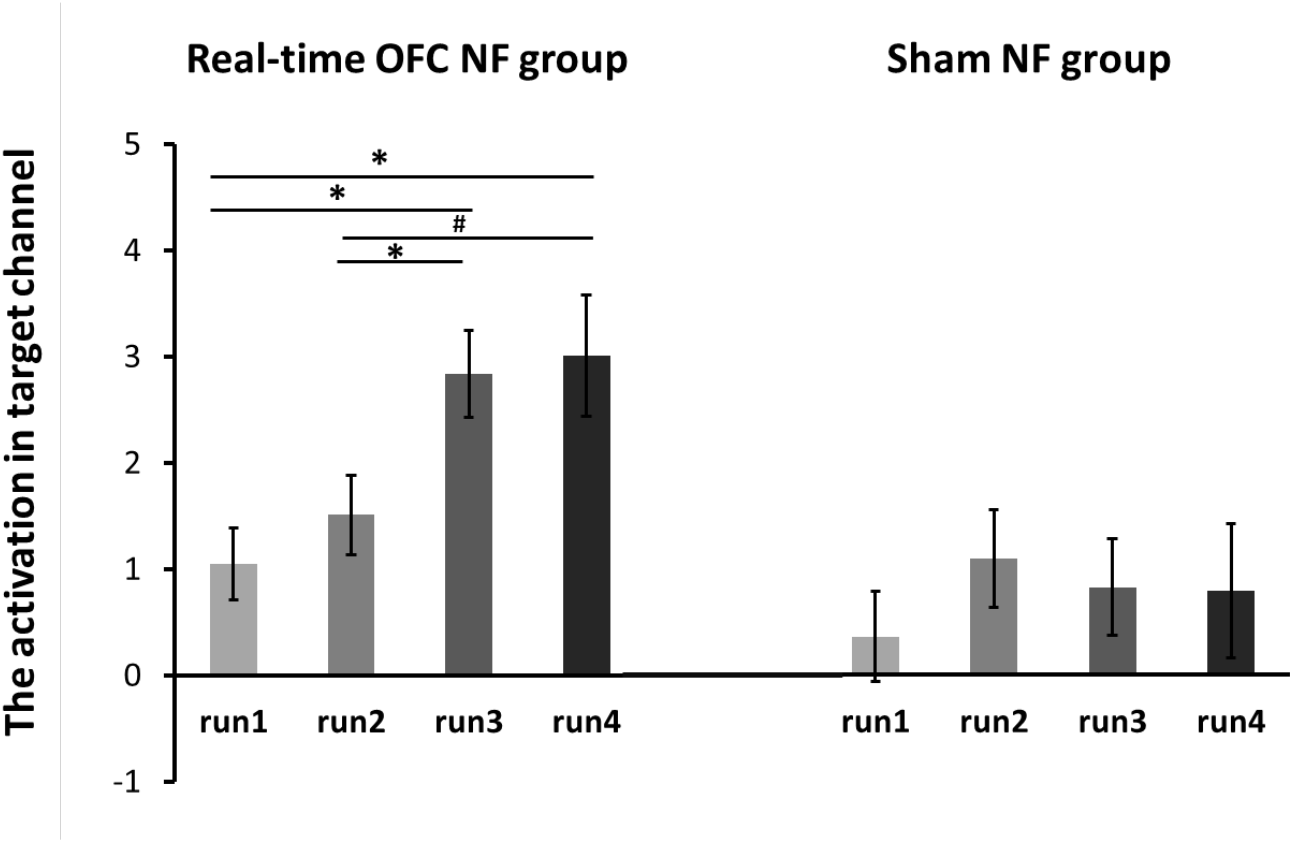
The activation in the target channel significantly increased over the course of the real-time OFC NF training runs but not during the sham NF training. Differences between the training runs were tested by post-hoc paired t-tests, two tailed. *p < 0.0125. # denotes marginal significance, p < 0.05.

### Exploratory analysis – regional specificity of the training effects

To explore the regional specificity of the neural training effects of training on all OFC channels was explored. A mixed ANOVA with the factors run (run1/run2/run3/run4), group (real feedback vs. sham feedback) and channel (channel No.1-No. 12) and the dependent variable OFC activity as measured by the beta-values revealed a main effect of run (F_(3,162)_ = 4.14, p = 0.013), a main effect of channel (F_(11,594)_ = 6.84, p < 0.001, degrees of freedom Greenhouse-Geisser adjusted), a run*group interaction effect (F_(3,162)_ = 2.93, p = 0.047) and a channel*group interaction effect (F_(11,594)_ = 4.49, p = 0.001), but no main effect of group and no other interaction effects. To control for multiple comparisons a Bonferroni correction was used to account for all channels tested. Post-hoc comparisons demonstrated that the significant differences between experimental and control group were observed in the target channel No.7 (p < 0.001, two-tailed) and the adjacent channel No.6 (p=0.012, two-tailed) in the lateral OFC (details see Figure 1 and Figure 4). For all other channels the interaction effect was not significant (all ps > 0.132, two-tailed), indicating that training specifically modulated activity in the right lateral OFC.

**Figure 4.**
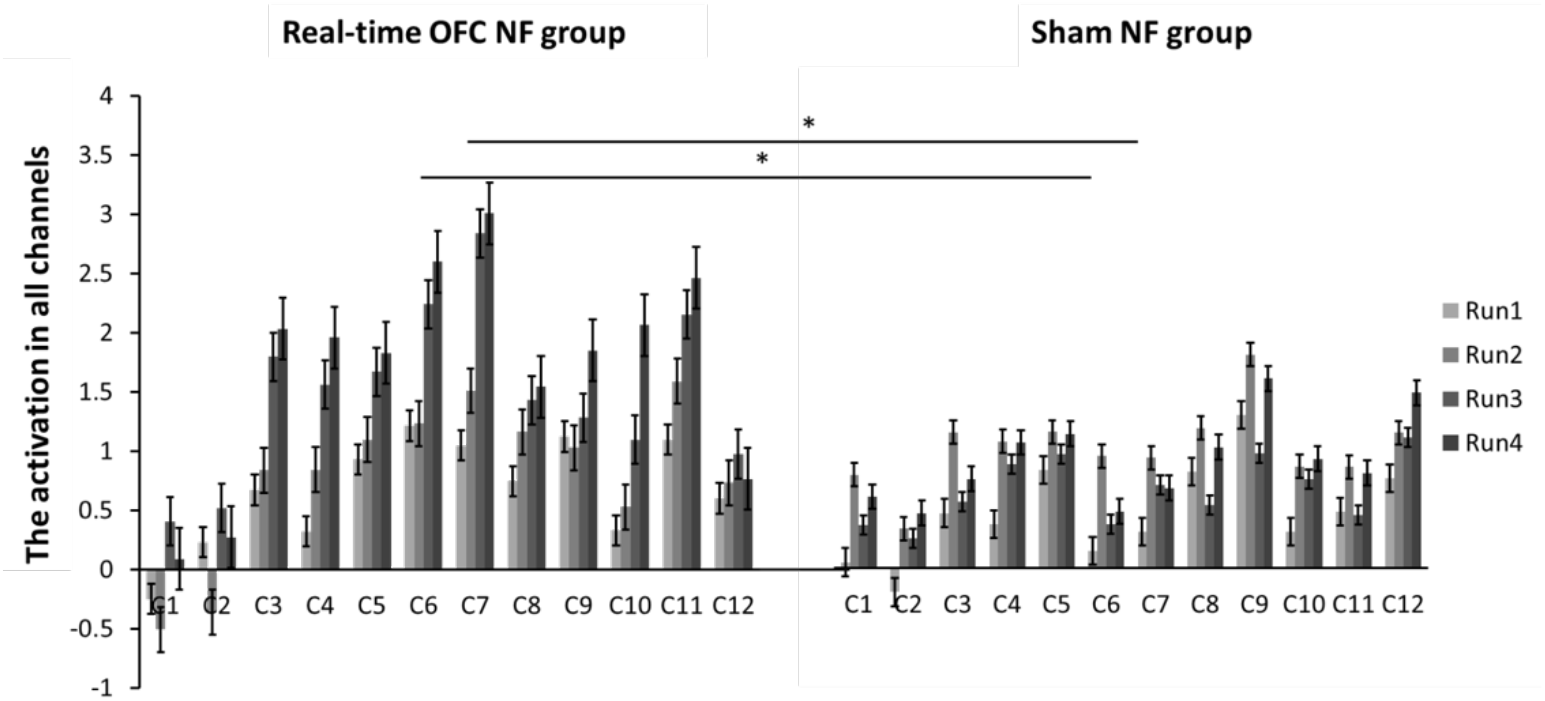
Significant differences between the experimental and sham control group were observed in the target channel 7(C7) and the adjacent channel 6 (C6) in the lateral OFC. Differences between groups were tested by means of post-hoc two sample t-tests, two tailed. *p < 0.004.

### Evaluation of training success – primary behavioural outcome

Performance indices for the IED task included the number of errors, the number of trials completed, the number of stages completed and reaction times. In line with previous research the median reaction times were used as estimate of central tendency [53]. Results revealed significant shorter reaction times for correct responses after the experimental training compared to the sham training (p = 0.037, one-tailed, two-sample t-test, see figure 5). Other indices of the IED task and rewarding experience failed to reach statistical significance (all p > 0.05, one-tailed).

**Figure 5.**
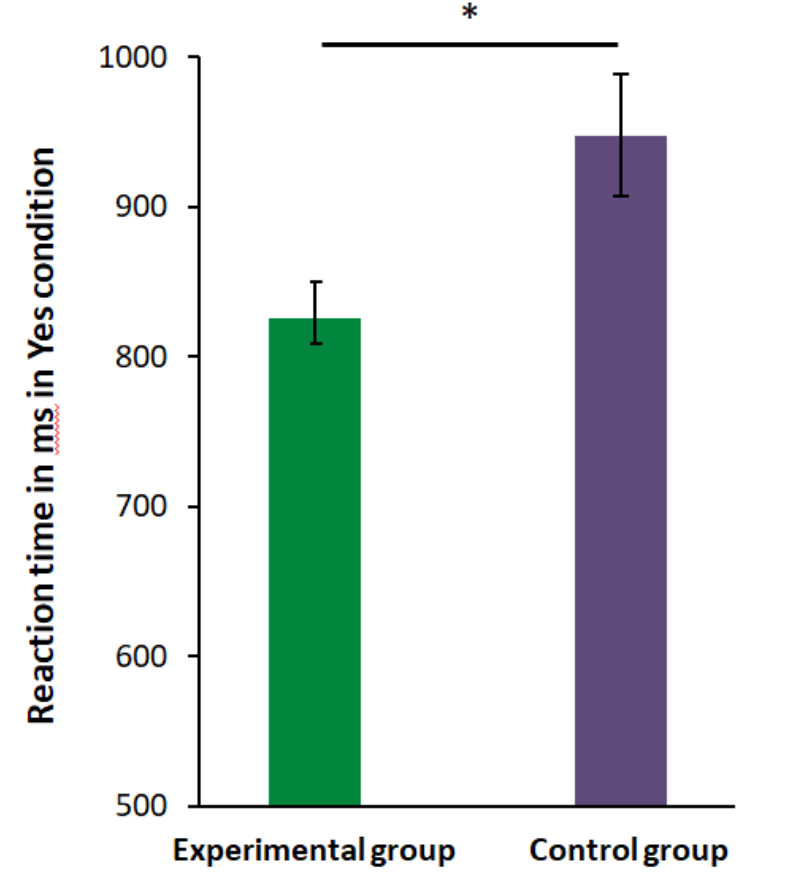
Significant differences between the experimental and control group were observed for reaction time for correct responses in the IED paradigm. Between-group differences were tested by means of two sample t-tests, one tailed. *p < 0.05.

### Association between behavioural and neural training success

A subsequent correlation analyses examined the association between behavioural (median reaction time in the IED task) and neural training success (changes run4 > run1 activation in target channel). Results revealed that reaction times for correct responses showed a descriptive, but not significant, negative association with OFC activity changes in the training group (r_experimental_ = −0.289, p = 0.144, see figure 6A) but a marginal positive association in the sham group (r_control_ = 0.361, p = 0.055; stable after excluding one outlier, r_control_ = 0.097, p = 0.622, significant correlation differences according to Fisher’s Z test, z = −2.41, p = 0.016; see figure 6A).

**Figure 6.**
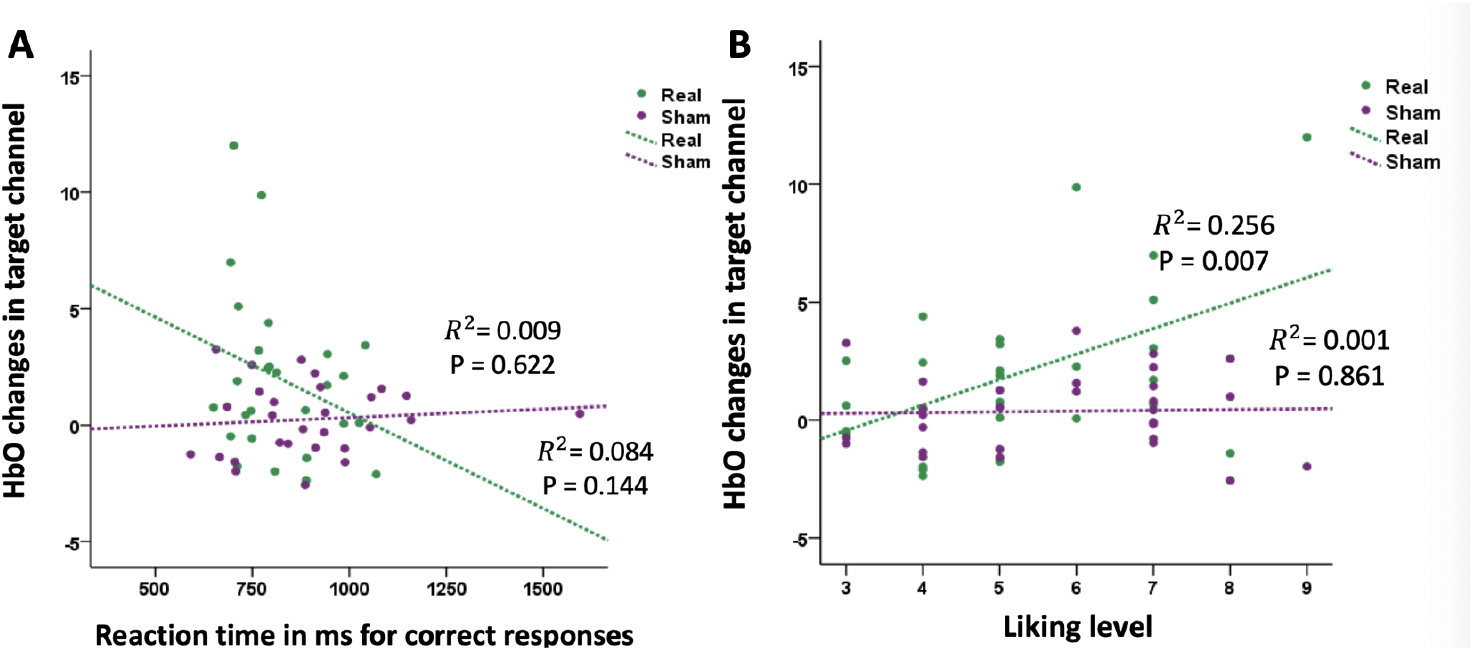
A. In the experimental group stronger training-induced OFC activity changes (run4>run1) were negatively associated with IED response times, whereas a positive association was observed. In the sham group. B. In the experimental group stronger OFC changes were positively associated with higher levels of liking, in the sham group no association was observed.

Given that a previous study reported significant associations between fNIRS-assessed OFC activity and liking levels [54], associations with post-training liking ratings were additionally explored. Results indicated that the liking level was significantly positively associated with OFC activation changes in the training (r_experimental_ =0.506, p = 0.007 see figure 6B), but not the sham control group (r_control_ = 0.034, p = 0.861, marginal significant between-group correlation differences, z = 1.87, p = 0.062; see figure 6B).

### Regulation strategies reported by the participants

In line with a previous study that evaluated regulation strategies during neurofeedback training with the present platform [60], the content analysis identified three main clusters of up-regulation strategies: (1) imagination (‘Imaging I will reach a greater level of career success’), (2) experience recall (‘Thinking about my happy memories’); (3) meditation (‘Thinking of nothing particular and relaxing”). Importantly, the groups did not differ in the regulation strategies employed during the training (Pearson c ^2^ test, p = 0.734, two-tailed,

Table 2), arguing against confounding effects of different regulation strategies on the observed neural and behavioural between-group differences.

**Table 2.**
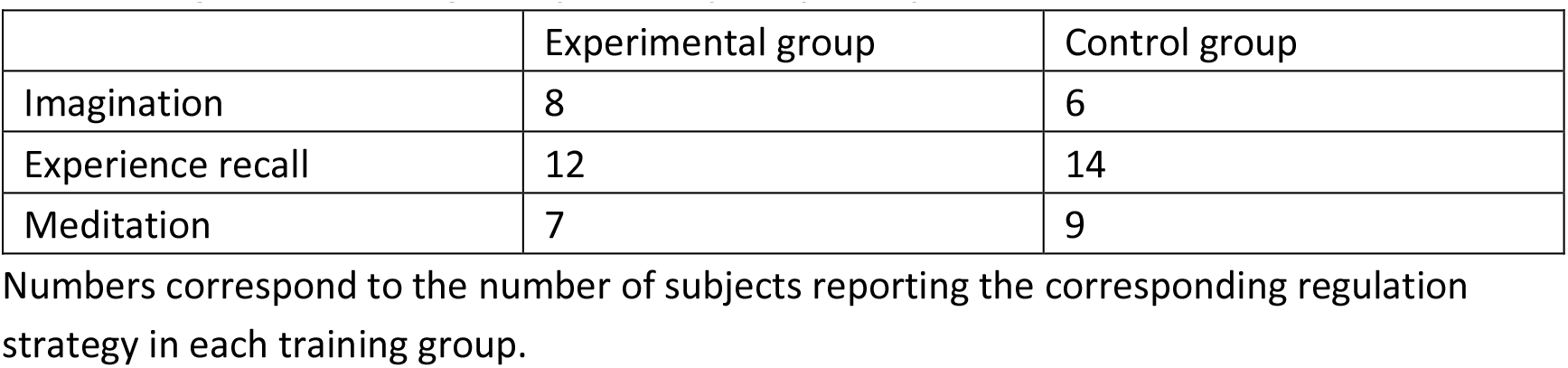
Regulation strategies reported by the participants.

## Discussion

The present proof-of-concept study evaluated the feasibility and functional relevance of real-time fNIRS-informed neurofeedback (NF) training as closed-loop strategy to increase lateral OFC (lOFC) activitation. Using a randomized sham-controlled between subject design the present study revealed that participants in the experimental group successfully learned to increase lOFC activity over the course of four subsequent training runs. Importantly, no significant changes in neural activation were observed in the sham control group. Together with the lack of between-group differences in perceived training success and regulation strategies the lack of training-induced changes in the sham group argues against unspecific effects of the training procedure and emphasizes the specific importance of the feedback signal for successful acquisition of neural regulation. Exploring the regional specificity of the training effects on OFC activation revealed that significant training-induced changes were restricted to the target and an adjacent channel suggesting that the training produced regional-specific increases in right lOFC activation. On the behavioral level the experimental group demonstrated a trend for enhanced cognitive flexibility as reflected by decreased response times for correct responses in the attentional set-shifting task. Moreover, exploratory analyses revealed that shorter response times and higher rewarding experience were associated with stronger training-induced increases in lOFC activity, further confirming a potential functional relevance of successful lOFC regulation via fNIRS-informed neurofeedback.

Comparing the experimental training with the sham control group demonstrated that fNIRS-informed neurofeedback from the target channel allowed subjects to acquire regulatory control over regional-specific activation in the OFC. Examining the activation changes within the groups further documented that lOFC activity significantly increased over the four training runs in the experimental but not the sham control group. The successful up-regulation in the training group was further confirmed by between group analyses showing significantly higher lOFC activity in the experimental group compared to the sham group during the last two training runs. The present results add to the growing number of reports suggesting that fNIRS-informed neurofeedback training allows subjects to gain volitional control over cortical brain activity, including motor as well as frontal regions [32-34]. Together with a previous study employing fMRI-informed neurofeedback to train subjects to volitionally modulate orbitofrontal activity [10], the present findings additionally document that hemodynamic neurofeedback-assisted control over regional activation in the OFC is feasible. Compared to fMRI-informed neurofeedback fNIRS-informed neurofeedback is limited by characteristics inherent to the acquisition methodology, particularly a lower spatial resolution and restricted signal-acquisition from cortical regions. However, particularly for clinical applications or for targeting regions susceptible to MRI-artifacts such as the OFC, the advantages of fNIRS-informed neurofeedback may outweigh the limitations and promote translation into the clinical practice. Orbitofrontal alterations have been demonstrated in several disorders characterized by deficits in cognitive flexibility and value processing, particularly attenuated OFC activation during impaired cognitive flexibility in paediatric and adult obsessive-compulsive disorder [55,56] and deficient value-guided response selection in substance addiction have been reported [57]. These regional alterations may reflect network-level dysfunctions in striato-frontal circuits, specifically the lateral OFC-caudate pathway engaged in attentional shifting and cognitive flexibility [15]. fNIRS-assisted regulatory control over the lOFC may promote normalization of aberrant neural activation and promote functional recovery in psychiatric populations.

Examining the training effects on behavioral domains associated with the lOFC revealed some evidences for the functional relevance of the training. Although no effects on accuracy in the set shifting task were observed, the experimental group demonstrated slightly faster reaction times for correct responses as compared to the sham control group. The OFC critically contributes to flexible behavioral adaptations, with the lateral region supporting adaptation to changing reward contingencies during reversal learning [16-20,58]. However, in the present study regional-specific modulation of the lOFC did not affect the acquisition of the dimensional shifts per se. Despite several human imaging studies suggesting a role of the OFC in cognitive flexibility [23,59], previous animal studies demonstrated that regional-specific lesions of the lateral OFC did not critically disrupt set-shifting performance which may explain the lack of strong performance effects on the set-shifting paradigm in the present study [20,60]. Despite a lack of between-group differences in the subjective liking of the task, an exploratory correlational analysis revealed that stronger training-induced lOFC increases in the experimental group associated with higher post-training liking ratings whereas no such association was observed in the sham group. These findings are in line with a previous report on a positive association between lOFC activation and subjective liking during a social reward paradigm [54] and the important role of the lOFC in processing reward-related outcome expectancies [61].

The present findings need to be interpreted in the context of limitations. First, to reduce variance related to sex-differences or effects of menstrual cycle on OFC-related functions and neural activity, the present proof-of-concept study focused on male participants (for a similar approach see [6]). The generalizability of the present results to female participants and potential sex-differences thus need to be examined in future studies. Secondly, despite some evidences for effects of lOFC neurofeedback training on the behavioral level, the between-group comparisons of the primary behavioral outcomes failed to reach statistical significance. The lack of robust effects may be explained in terms of a low sensitivity of the set-shifting paradigm for activation changes in the lateral OFC. Moreover, the present study employed a single training session and more intense training schedules may be necessary to produce robust behavioral effects. Finally, previous studies using fMRI-informed neurofeedback recently demonstrated that participants can maintain regulatory control for several days and in the absence of feedback [4,6]. The present proof-of-concept study did not include a follow-up or maintenance session and thus the maintenance of fNIRS-assisted lOFC regulatory control and its independence of online feedback remain to be determined in future studies.

In summary, the present findings demonstrate that real-time fNIRS-informed neurofeedback training of the OFC is feasible and that this approach allows subjects to volitionally increase activation in this region. Given the high clinical relevance of altered OFC activity in psychiatric disorders, fNIRS-informed training of this region may represent a promising strategy to normalize OFC function in these disorders.

## Acknowledgements

The authors declare no competing interests.

